# Identification of immune response genes of cassava during early phases of cassava brown streak virus infection

**DOI:** 10.1101/2025.04.29.651173

**Authors:** Jessica Lilienthal, Stephan Winter, Boas Pucker, Samar Sheat

## Abstract

Cassava brown streak disease (CBSD) caused by the ipomoviruses CBSV and UCBSV, poses a significant threat to cassava crops and the destruction of tuberous roots leads to substantial yield losses. Virus resistance identified in some South American cassava germplasm lines inhibits virus replication and movement and, is a complete immunity of some lines or, a restriction of the virus replication to phloem companion cells and root infection in others. To further explore the resistance responses and identify potential resistance genes, a time-course RNA-Seq study was conducted comparing gene expression of a virus-susceptible cassava TMS 96/0304 with a cassava line restricting CBSV to the roots and a cassava DSC 167 (COL2182), immune to the virus. Differential gene expression patterns across leaves, stems, and roots revealed substantial differences between susceptible and resistant genotypes. Temporal analysis highlighted significant variations in the number of differentially expressed genes (DEGs), particularly in stem tissues at one day after inoculation (DAI), with notable overexpression observed in the resistant lines DSC 167 (436 DEGs) and DSC 260 (335 DEGs). In contrast, significant expression changes were noted only at 10 DAI in the virus infected TMS 96/0304 (737 DEGs). Several DEGs in resistant genotypes were associated with proteins involved in hormonal regulation, defense and immunity, transcriptional regulation, stress response, cell structure, and signal transduction. The analysis indicates that virus resistance in the studied lines is driven by early-stage regulation of genes for which specific functions remain to be elucidated.

## 1. Introduction

Cassava (*Manihot esculenta* Crantz) is a woody shrub from the family *Euphorbiaceae* and is cultivated in South America, Africa, and Asia (Thresh 2006; Alves 2002). It is a crucial food security crop in Africa, providing the nutritional foundation for many people (Thresh 2006; Howeler *et al*. 2013). In Africa, cassava is threatened by several diseases and the cassava brown streak disease (CBSD) is having the most severe impact. Two distinct virus species cause CBSD; the cassava brown streak virus (CBSV) and the Ugandan cassava brown streak virus (UCBSV). They belong to the family *Potyviridae* and the genus *Ipomovirus*, (Winter *et al*. 2010; Monger *et al*. 2001) have distinct single-stranded RNA genomes but induce similar symptoms in infected cassava (Winter *et al*. 2010; Munganyinka *et al*. 2018; Hillocks and Jennings 2003). Brown lesions appear on the stem, leaves exhibiting vein clearing and chlorosis (Hillocks *et al*. 2002) and tuberous roots, dark brown, necrotic areas often throughout the entire tuber are prominent signs of the diseases (Nichols 1950; Patil *et al*. 2015). Root necrosis can lead to a complete loss of the crop, presenting a significant threat to cassava production and harvest (Alicai *et al*. 2007; Story, H. H. 1936). The viruses are transmitted by whiteflies (*Bemisia tabaci*) in a semi-persistent manner but the disease mainly spreads through the exchange of infected stem cuttings used for vegetative propagation (Story, H. H. 1936; Maruthi *et al*. 2005).

Extensive efforts to develop CBSD resistant cassava lines only reached limited success because resistant varieties among African cassava were not available. Novel sources of virus resistance thus were identified in South American germplasm accessions (Sheat *et al*. 2019; Sheat 2020) and the most promising lines were subsequently incorporated into breeding programs to create cassava with resistance against viruses causing CBSD and against viruses causing cassava mosaic disease (CMD) which is endemic wherever cassava is grown on the continent (Sheat *et al*. 2022; Sheat and Winter 2023). Highly stringent methods were used to screen for strong and durable resistance lines. Detailed studies on the resistance phenotypes (Sheat *et al*. 2019) revealed two distinct resistance categories: highly resistant (immune) plants, such as DSC 167, free of symptoms (Kaweesi *et al*. 2014) and with virus not detectable and plants, such as DSC 260, with virus invasions and symptoms confined to roots while the rest of the plant remained virus-free. Further studies (Sheat 2020) showed that the resistance was associated with the restriction of the virus to the companion cells of the phloem. Interestingly, in the immune line DSC 167, CBSV was restricted to the external phloem tissues while external and internal phloem was invaded in DSC 260 (Munganyinka *et al*. 2018; Sheat 2020). However, a mechanistic explanation for this phenomenon is still pending.

RNA-seq transcriptome sequencing is fundamental to most analyses of molecular processes regulating gene expression. Comparative studies of resistant and susceptible cassava were done using RNA-seq to analyze virus infections and cassava resistance response 28 days after introducing a mixed infection of CBSV and UCBSV (Anjanappa *et al*. 2018). In a study by Maruthi *et al*. 2014, RNA-seq was used to compare susceptible and resistant cassava varieties to find genes involved in resistance against CBSV and UCBSV. The author showed that the resistant varieties had a high number of overexpressed genes among which NAC genes from hormone signaling and phenylpropanoid pathway were found most significant (Maruthi *et al*. 2014). In the research of Amuge *et al*. 2017, transcriptome analyses were conducted at early time points after graft-infection (6 hours after grafting) and 1, 2, 5, and 8 days thereafter. At 2 and 5 days after infection, many plant defense genes were differentially expressed in the resistant genotype, including translation elongation factors, NBS-LRR, heat shock proteins, and pathogenesis-related proteins (Amuge *et al*. 2017). The study suggested that even earlier time points after infection should be analyzed (Anjanappa *et al*. 2018; Amuge *et al*. 2017).

We questioned the virus resistance character of the cassava lines used in earlier reports (Sheat *et al*. 2019) and based our RNA-seq analysis on new sources of resistance in cassava lines that were thoroughly tested over many years for their response against viruses causing CBSD. The three cassava lines – two resistant South American germplasm accessions (DSC 167 and DSC 260) and a susceptible line (TMS 96/0304) were subjected to RNA-seq analysis at 1, 5, and 10 DAI. Using transcript abundances as a proxy, gene expression at the defined time points was analyzed to determine differential gene expression highlighting genes activated during an initial response to infection and genes involved in regulating viral replication and movement in early immune response of cassava against CBSV.

## 2. Results

### 2.1. Symptom Expression in CBSV-Infected Cassava Plants

Except for the susceptible TMS 96/0304, the resistant cassava DSC 260 and DSC 167 at no time point after graft-infection, did not show symptoms of CBSD infection on leaves and stems. One month after grafting CBSV infections were clearly visible on stems of infected TMS 96/0304 plants manifesting as brown lesions near the infection site. Two of the remaining plants exhibited symptoms on the stem and leaves scoring severe symptoms (S++) while the other remaining four plants had mild to moderate symptoms on stems (S+).

### 2.2. Read Mapping Against the Cassava Brown Streak Virus Genome

To trace virus replication, the trimmed reads from RNA-Seq were aligned to the CBSV-Mo83 genome sequence (FN434436.1, NCBI). No viral reads were found in RNA-Seq of all tissues at 1 and 5 DAI (Table 1). At 10 DAI, viral reads were identified in stem and root tissues of TMS 96/0304 and DSC 167 (Table 1) while DSC 260 remained virus-free. In TMS 96/0304, 2,001 and 970 reads were detected to replicate A and B respectively while a considerably higher number of viral reads was recorded in samples from roots with 14,894 and 17,614 in replicate A and B, respectively. No viral reads were detected in the root-restricted line DSC 260 while a stem and root sample of the resistant cassava DSC 167 contained many viral reads (Table 1). The entire read mapping numbers and percentages are listed in Table S1-S4.

**Table 1.**
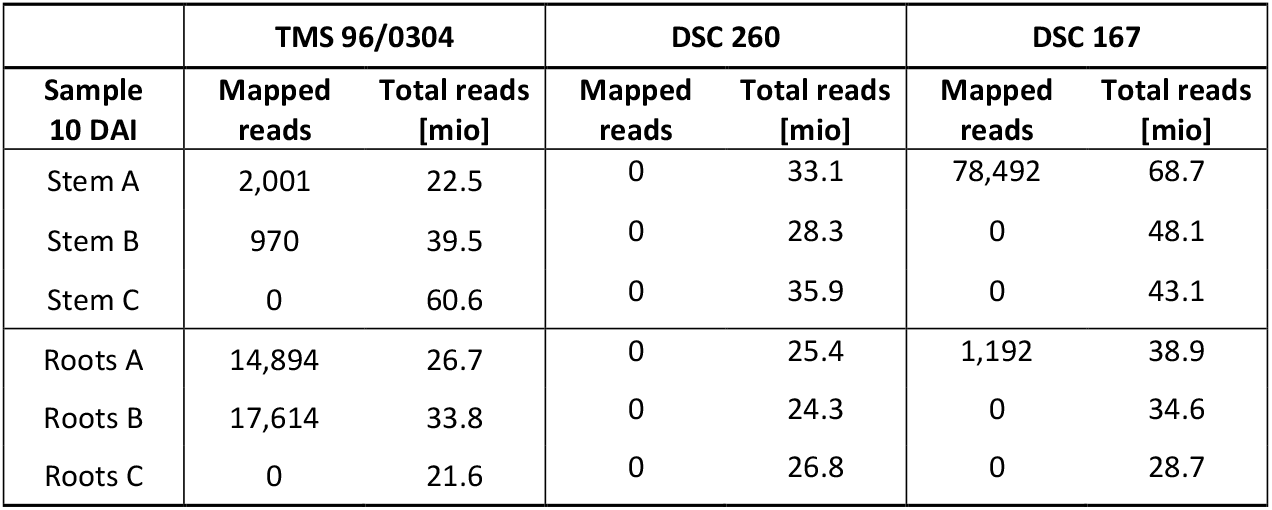
Viral reads recorded 10 DAI in infected stem and root tissue of three resistant and susceptible cassava lines.

|  | TMS 96/0304 |  | DSC 260 |  | DSC 167 |  |
| --- | --- | --- | --- | --- | --- | --- |
| Sample 10 DAI | Mapped reads | Total reads [mio] | Mapped reads | Total reads [mio] | Mapped reads | Total reads [mio] |
| Stem A | 2,001 | 22.5 | 0 | 33.1 | 78,492 | 68.7 |
| Stem B | 970 | 39.5 | 0 | 28.3 | 0 | 48.1 |
| Stem C | 0 | 60.6 | 0 | 35.9 | 0 | 43.1 |
| Roots A | 14,894 | 26.7 | 0 | 25.4 | 1,192 | 38.9 |
| Roots B | 17,614 | 33.8 | 0 | 24.3 | 0 | 34.6 |
| Roots C | 0 | 21.6 | 0 | 26.8 | 0 | 28.7 |

### 2.3 Clustering of Samples by Tissue Type

Principal component analysis (PCA) of the TPM (transcripts per million) values classified each sample according to its specific tissue type (Figure 1). The stem, root, young leaf (YL), and old leaf (OL) samples grouped within their respective regions, demonstrating strong coherence among biological replicates. Six outliers were identified in the root samples which is likely due to low values detected during quality control assessments using Qubit and Bioanalyzer. The outliers recorded for YL and stem samples were attributed to biological variability since similar rRNA content was also recorded.

**Figure 1.**
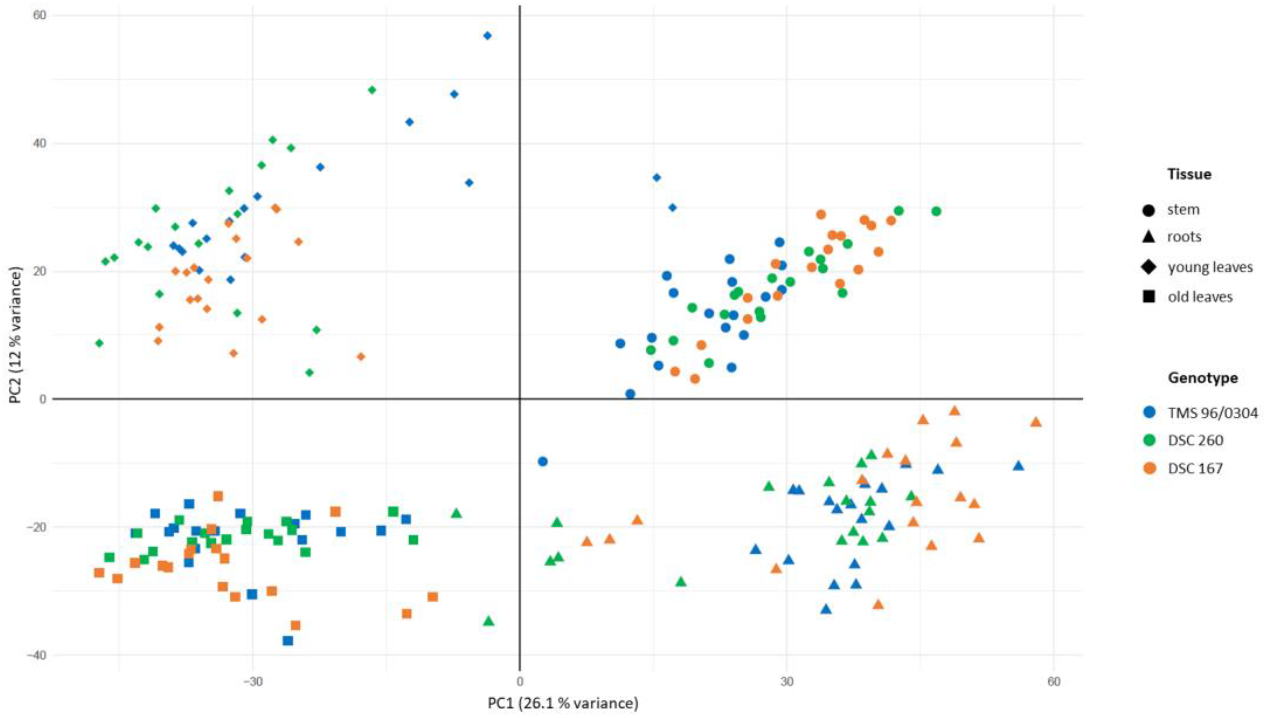
PCA plot of the TPM values of all samples, OL (blue), YL (green), roots (red), stem (brown). Variance of PC1 by 26.1 % and PC2 by 12 %. Distribution of the samples by the type of tissue. **(A)** Young leaves, YL; **(B)** Stem tissue; **(C)** Old leaves, OL; **(D)** Roots.

Further PCA plots specific for each tissue type revealed a read distribution according to cassava genotype (Figure S1). The PCA plot of all stem tissue samples discriminated reads of the susceptible TMS 96/0304 from those of two resistant lines showing some overlap. A similar PCA pattern was evident in the root and YL samples, where genotypes separated into distinct clusters within only minimal overlap. In contrast, the PCA plot of the OL samples displayed a markedly different pattern, where samples could not be explicitly classified into three groups.

### 2.4 Gene Expression in Virus-Infected Cassava 1, 5, and 10 Days After Virus Inoculation

Upon graft infection with CBSV, cassava genotypes with virus resistance (DSC 167, DSC 260) showed a rapid response to virus infection already at 1 DAI which was characterized by a steep increase of gene expression in stem, roots, YL, and OL (Figure 2A). At this time point, there was no response from the susceptible TMS 96/0304 and the mock-grafted control group. At 10 DAI, the number of DEGs in the susceptible TMS 96/0304 increased strongly while the number of DEGs in the resistant genotypes, DSC 260, and DSC 167, significantly decreased (Figure 2C). Interestingly the time point 5 DAI remained without findings and considerable gene expression differences were only found in the OL of the highly resistant cassava DSC 167 (Figure 2B). Since the distribution of OL reads in the PCA plot and their number of DEGs in the bar charts were not informative, RNA-Seq data from OL were not further analyzed. The total number of differentially expressed genes (DEGs) identified is presented in Table S5.

**Figure 2.**
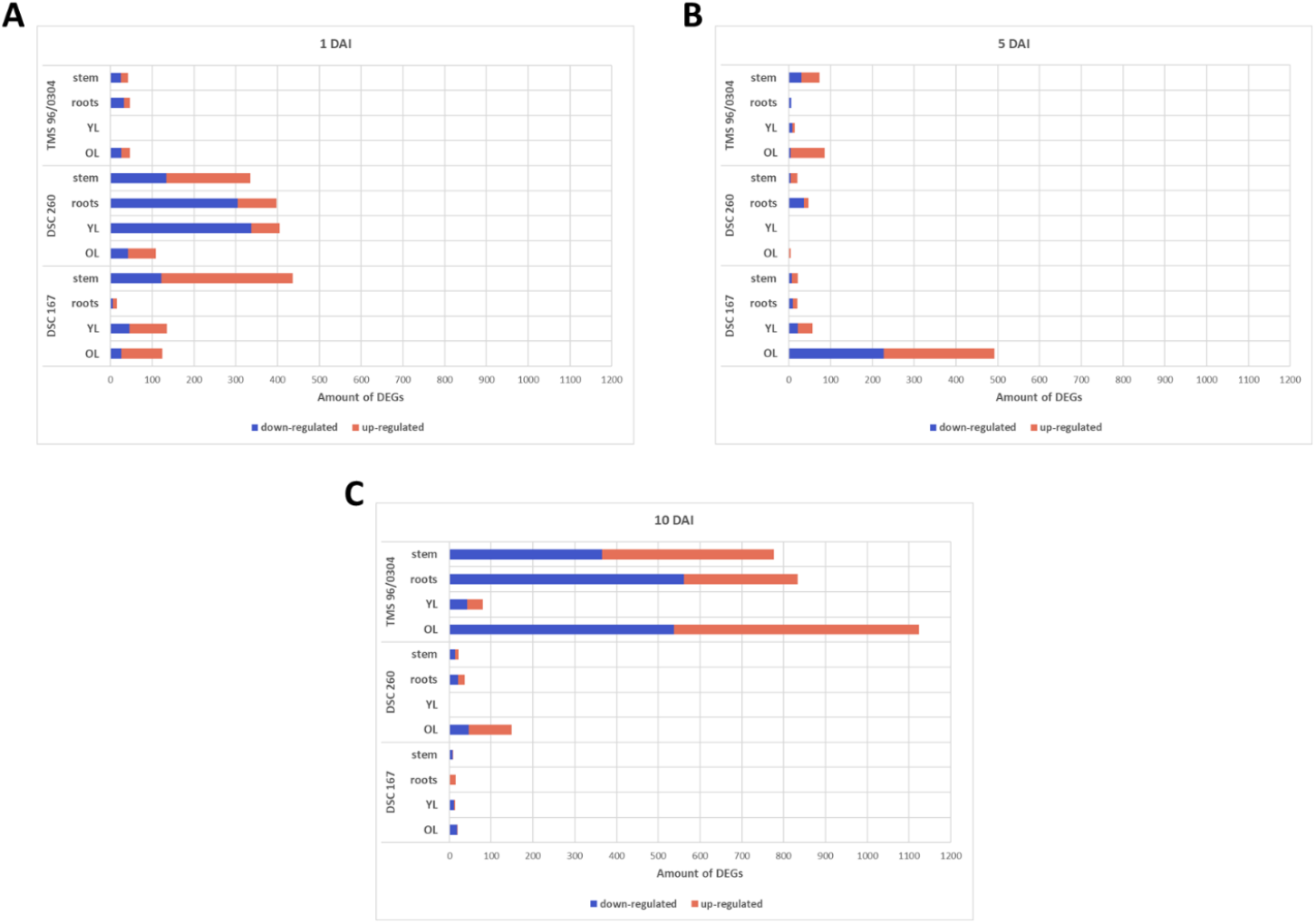
Bar charts of DEGs for each genotype and tissue. Thresholds for adjusted p-value < 0.05 and |log2 fold change| > 1. Up-regulated genes (orange), down-regulated genes (blue). **(A)** 1 DAI; a high number of DEGs in root-restricted DSC 260 and resistant DSC 167 cassava plants. **(B)** 5 DAI; no significant DEG except in OL of the resistant genotype DSC 167. **(C)** 10 DAI; significant DEG in the susceptible genotype TMS 96/0304 while no changes were recorded for resistant cassava lines.

### 2.5 Genes Involved in an Early Plant Response against CBSV Infection

Protein IDs of the DEGs were mapped for functional annotation to entries using NCBI (accessed in September 2024). The resulting genes were allocated to distinct gene clusters associated with hormonal regulation, transcriptional regulation, stress response, defense and immunity, cell structure, and signal transduction. A total of 24 gene categories were formed within these gene clusters, and the functions of the individual genes within each category were identified. The complete list of all genes, proteins, their respective functions, regulation, and gene groups, is presented in Table S1.

For each DAI, genotype and tissue type, the number of DEGs belonging to a given gene category was counted. The resulting heatmap (Figure 3) illustrates the net trend within each group, calculated as the difference between the number of up- and down-regulated genes (e.g. -2 indicating two more down-regulated than up-regulated genes). This approach emphasizes overall regulatory patterns rather than absolute expression levels and does not account for the magnitude of differential expression.

**Figure 3.**
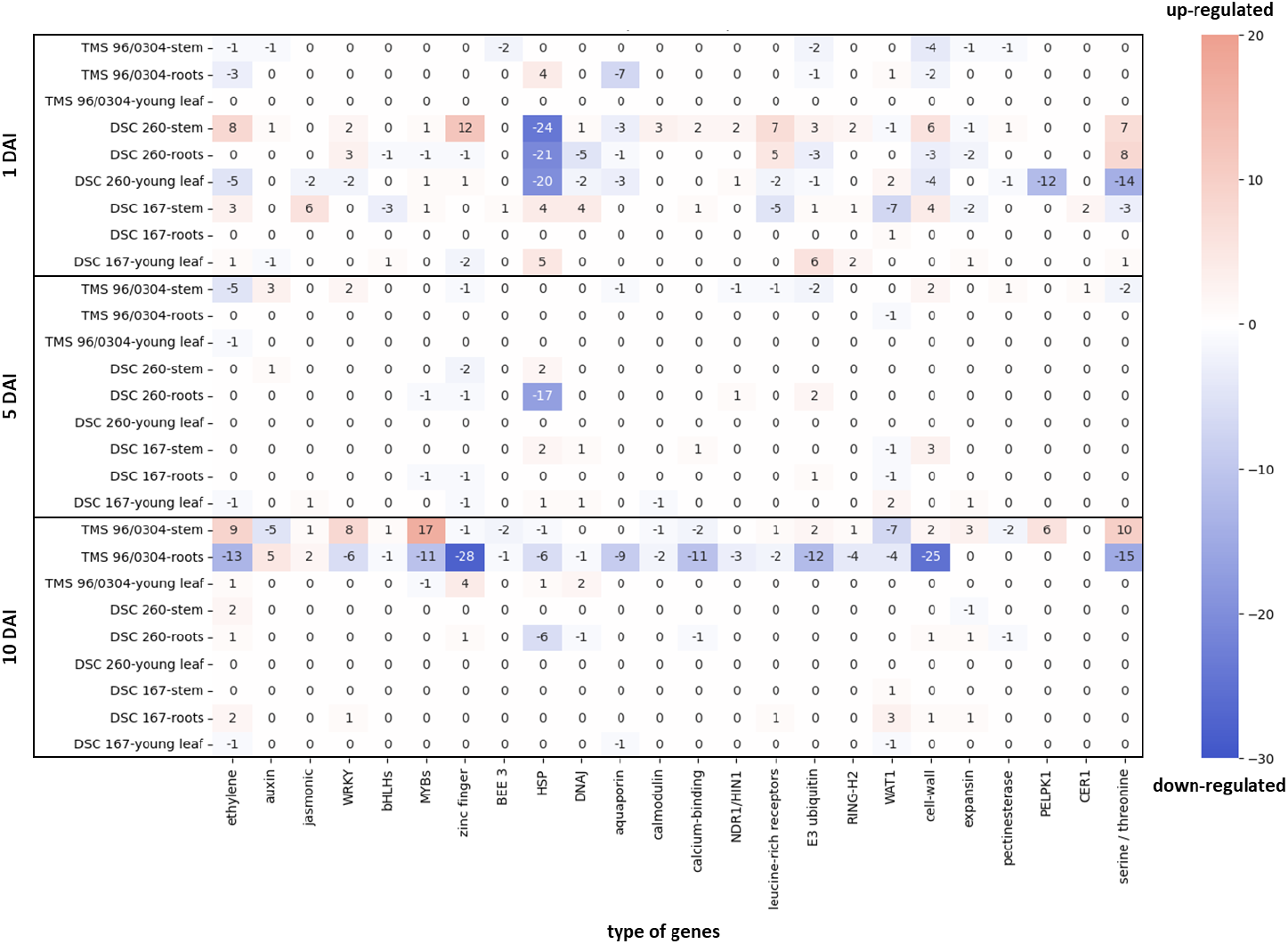
Heatmap of DEGs based on DAI, genotypes, and tissue types plotted against plant immune response genes. Significant down-regulation of almost all gene categories at 10 DAI in the virus infected TMS 96/0304 at 10 DAI. Strong up-regulation in the stem tissue of the resistant genotypes DSC 260 and DSC 167 1 DAI and in the TMS 96/0304 10 DAI. Threshold for adjusted p-value < 0.05 and |log2 fold change| > 1. Down-regulation (blue), up-regulation (orange).

Upon CBSV infection, plant immune response genes showed notable alterations already at 1 DAI in the resistant DSC 167 and DSC 260 cassava. In DSC 167 an increased gene expression of ethylene, jasmonic acid, HSPs, DnaJ and cell wall biosynthesis was observed in stem tissues and an HSP and E3 ubiquitin increase in leaves while there were no significant gene expression changes in roots. In contrast, DSC 260 showed and increased level of ethylene, zinc finger, calmodulin, leucine rich receptors, cell wall biosynthesis and serine/ threonine kinases in stem tissue while a dramatic downregulation of HSP was recorded for the roots all other tissues analyzed (Figure 3). The transcriptional responses of HSP genes was in sharp contrast to the upregulation of HSP in stem and leaf tissues. At 1 DAI, there were no differential expression recorded for roots of DS 167 and the susceptible TMS 96/0304 while critical changes were recorded for DSC 260. The most critical gene expression changes for the susceptible TMS 96/0304 were recorded only at 10 DAI in root tissues in which nearly all genes associated with plant immune responses were down-regulated. In contrast, RNA-Seq of stem tissues showed a notable up-regulation of genes involved in ethylene biosynthesis, WRKY, MYB, cell-wall, pectin esterase, CER1, and serine/threonine pathways. In contrast, RNA-Seq of stem tissues showed a notable up-regulation of genes involved in ethylene biosynthesis, WRKY, MYB, cell-wall, pectin esterase, CER1, and serine/threonine pathways.

Pathway enrichment analyses conducted on DEGs at 1 DAI and 10 DAI revealed six enriched KEGG pathways, phenylpropanoid biosynthesis (ath00940), protein processing in endoplasmic reticulum (ath04141), plant hormone signal transduction (ath04075), and plant-pathogen interaction (ath04626). Within these pathways, multiple gene groups were differentially expressed with HSP90, HSP70, HSP40, small heat shock factors (sHSFs), AUX/IAA, and cassava calmodulin-like proteins (CaM/CML) showing the most pronounced expression patterns.

Heat shock proteins HSP90, HSP70, HSP40 and sHSFs are implicated in protein processing in the endoplasmic reticulum, ER. In the resistant DSC 167 1 DAI, these genes were up-regulated in stem and YL tissues but remained unaffected in root tissues. In contrast, HSP genes were down-regulated in all tissues of DSC 260. At 10 DAI HSP90 was upregulated in the susceptible and HSP40 and sHSF were down-regulated. HSP70, HSP40, and sHSF were down-regulated in the roots.

AUX/IAA proteins of the plant hormone signal transduction pathway were differentially expressed in the stem and root tissues. This gene group was down-regulated at 10 DAI in the susceptible genotype in the stem and root tissue. In the resistant genotype, the AUX/IAA genes were up-regulated at 1 DAI in the stem and the root-restricted line DSC 260 started to up-regulate these genes at 5 DAI in the stem.

CaM/CML proteins (calmodulin/calmodulin-like proteins) belong to the plant-pathogen interaction pathway. In stem and root tissues of the resistant cassava genotypes. CaM/CML genes were up-regulated at 1 DAI. At 10 DAI, CaM/CML genes were down-regulated in TMS 96/0304 and DSC 260 1 DAI, however in the resistant line, these genes were not differentially expressed. The distribution of the gene regulation for each genotype, DAI and tissue type is shown in Figure 4.

**Figure 4.**
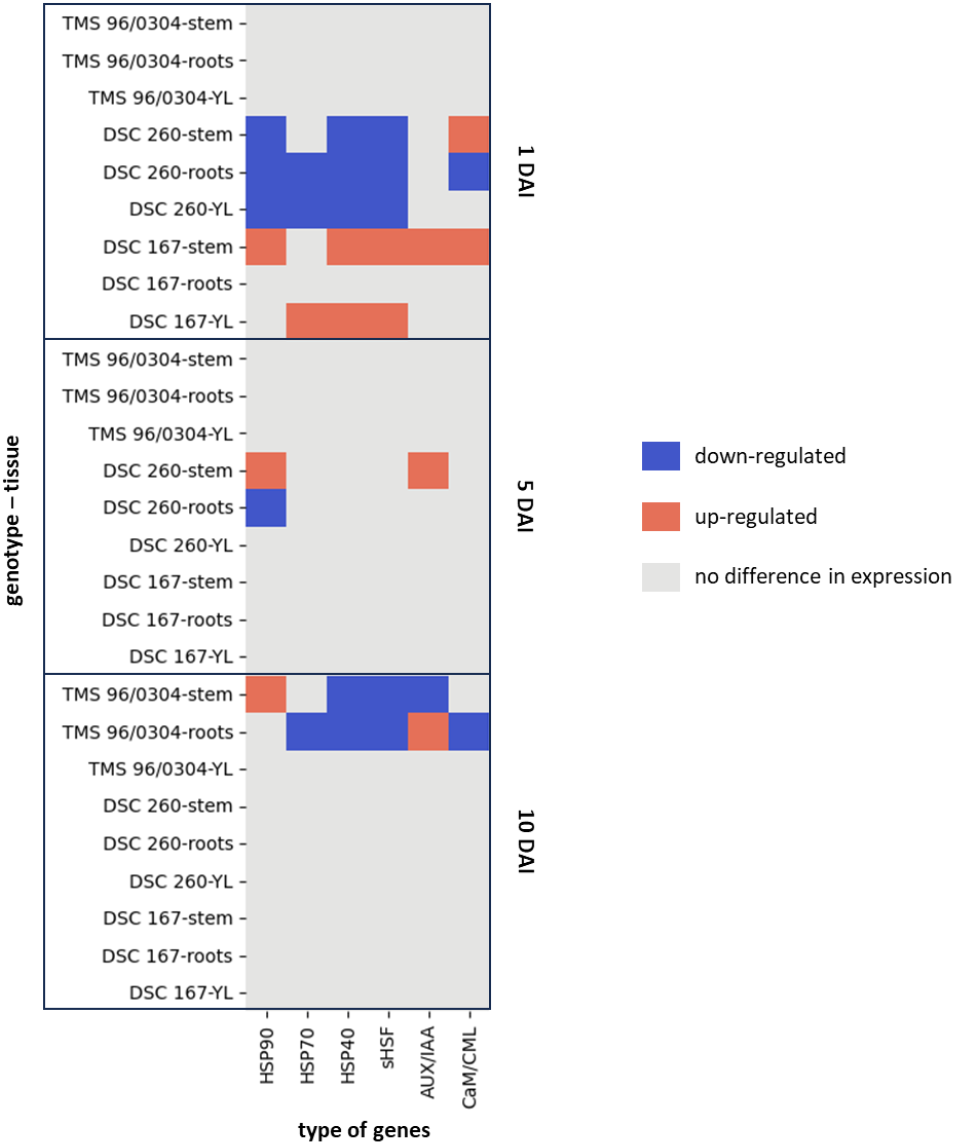
Heatmap of gene regulation for the three genotypes, tissue types and DAI plotted against the six gene candidates. Down-regulation (blue), up-regulation (orange), no difference in gene expression (grey). Significant down-regulation of the HSPs in the stem, roots and YL at 1 DAI in DSC 260 and up-regulation of genes in tissue of the stem and young leaves of the immune cassava line DSC 167. Strong differential response of the susceptible TMS 96/0304 involves downregulation of genes in roots and stem tissues.

Within these gene groups multiple genes were represented in the different genotypes (Table 2). For HSP90, the heat shock protein 83 was found while for HSP70 the heat shock cognate 70 kDa protein 2 was identified. The DnaJ protein homolog was represented by the HSP40 in this study. Multiple sHSF-encoding genes were found in the genotypes and tissues; however, the most frequent gene was the 23.6 kDa heat shock protein. For the AUX/IAA genes three proteins of the auxin-responsive proteins were found: IAA9, IAA14, and IAA27. The CaM/CML genes were associated with multiple different calcium-binding proteins: CML23, CML41, and CML43 (Table 2). The fold changes and gene IDs are shown in Table S5.

**Table 2.**
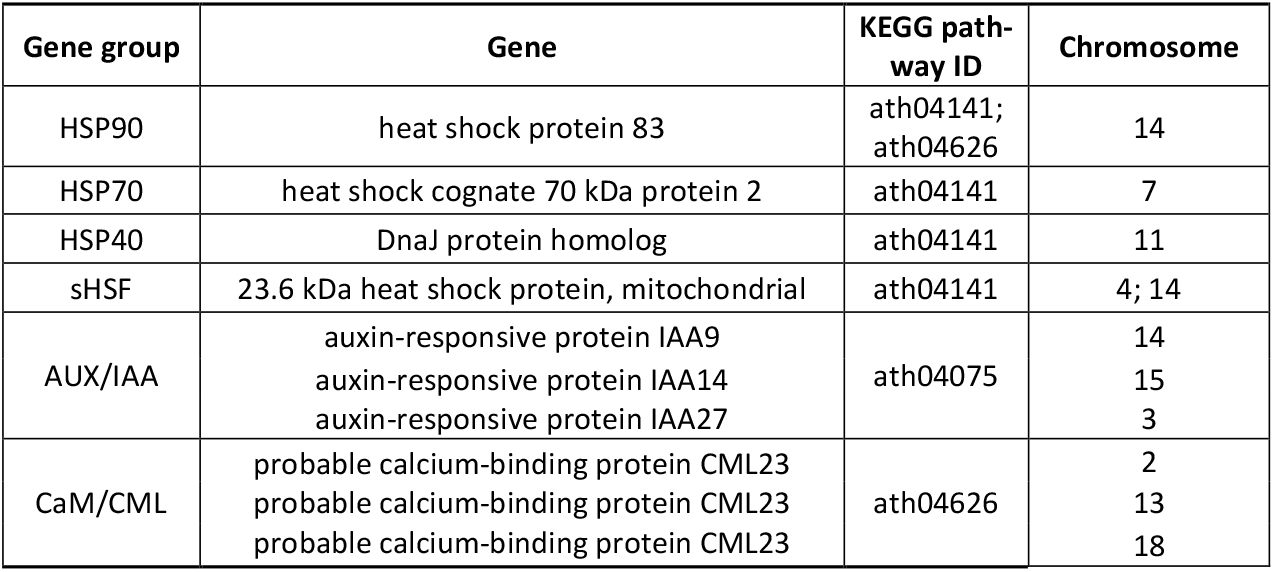
Gene candidates within their associated gene group and the KEGG pathways.

## 3. Discussion

This study employed RNA-Seq to investigate the transcriptomic responses of South American cassava genotypes exhibiting contrasting resistance to cassava brown streak viruses (CBSVs). By profiling gene expression at multiple time points post-inoculation, we aimed to uncover early molecular mechanisms that underpin resistance or susceptibility to CBSV. Our results provide a comprehensive overview of tissue-specific and genotype-dependent transcriptional reprogramming, highlighting key defense-related pathways activated in resistant cultivars.

Earlier transcriptomic studies have focused on African cassava genotypes such as Namikonga, Kaleso, and KBH2006/18 (Amuge *et al*. 2017; Maruthi *et al*. 2014; Anjanappa *et al*. 2016; Anjanappa *et al*. 2018) with contrasting response against infections with UCBSV and CBSV. The cassava materials chosen for our study were selected from cassava germplasm from South America that had been rigorously tested for its response to virus infections to select for immunity (DSC 167) and tissue specific resistance (DSC 260). In this resistance screening (Sheat *et al*. 2019), the cassava lines stated above failed our resistance tests.

One of the main challenges in cassava virus research lies in the difficulty of achieving uniform infection and defining a precise infection onset. These challenges introduce variability in controlled experiments and complicate interpretation. Nonetheless, our multi-timepoint RNA-Seq analysis revealed striking genotype-specific differences in the timing and magnitude of immune responses. Principal component analysis (PCA) indicated clear clustering by tissue type and genotype, with resistant genotypes showing marked transcriptomic shifts as early as 1 day after inoculation (DAI), while susceptible TMS 96/0304 displayed delayed responses, particularly at 10 DAI.

The immune genotype DSC 167 showed robust early induction of canonical defense-related genes, including heat shock proteins (HSP90, HSP70, HSP40), DnaJ, hormone signaling components (ethylene, jasmonic acid, and auxin-related genes), leucine-rich repeat (LRR) receptors, and genes associated with phenylpropanoid biosynthesis and cell wall reinforcement (Jones and Dangl 2006; Dodds and Rathjen 2010). Notably, these responses were strongest in stem tissues—identified here as the primary site of viral entry—underscoring the importance of localized early defense activation in preventing systemic virus spread. Conversely, both DSC 260 and TMS 96/0304 exhibited pronounced down-regulation of immune genes in root tissues at later time points. Down-regulated genes included HSPs, E3 ubiquitin ligases, aquaporins, and zinc finger transcription factors, correlating with increased viral loads. This repression of immune responses likely permits viral accumulation and systemic movement, especially in susceptible genotypes. These findings reinforce the concept that timing and tissue localization of defense activation are critical determinants of resistance.

HSP90 and HSP70 emerged as central players in the antiviral response. These molecular chaperones, along with DnaJ co-chaperones, are known to form complexes involved in protein folding, receptor stabilization, and immune signaling (Kanzaki *et al*. 2003; Tompa and Csermely 2004). HSP90’s role in stabilizing NB-LRR immune receptors is well-documented in other plant-pathogen systems, including resistance to Tobacco Mosaic Virus (TMV) in *Nicotiana benthamiana* and *Arabidopsis thaliana* (Hubert *et al*. 2003; Chen *et al*. 2010; Shirasu 2009; Takabatake *et al*. 2007). In our study, strong early up-regulation of HSP90 in stem tissue of DSC 167 suggests its involvement in rapid immune receptor activation and signaling. The differential expression of these chaperones in DSC 260 and TMS 96/0304 further supports their association with effective virus restriction.

AUX/IAA and CaM/CML genes, associated with hormone signaling and calcium-mediated stress responses respectively, were also up-regulated early in resistant genotypes. AUX/IAA proteins are key regulators of auxin signaling, and their interaction with salicylic acid and jasmonic acid pathways (Zhao *et al*. 2024) suggests a complex hormonal crosstalk during CBSV infection. Similarly, CaM/CML proteins, which mediate calcium signaling, may function as integrators of abiotic stress signals (Hu *et al*. 2018). The down-regulation of these genes in roots of susceptible genotypes points to a potential mechanism by which virus evades host defense by suppressing early signaling cues.

Together, these findings point to a model in which effective CBSV resistance in cassava depends on rapid and tissue-specific activation of immune pathways. The stem plays a central role as a surveillance hub, where early transcriptional activation of hormone-regulated and structural defense genes limits virus replication and movement. In contrast, delayed immune responses in the roots of susceptible lines enable systemic virus establishment.

To confirm the functional relevance of candidate genes such as HSP90, AUX/IAA, and CaM/CML, targeted knockdown or overexpression studies (e.g. via RNAi or CRISPR/Cas9) should be conducted. Elucidating their mechanistic roles in virus resistance will not only advance our understanding of cassava immunity but also inform breeding strategies aimed at developing resistant cultivars.

This work provides novel insights into early immune responses in South American cassava genotypes challenged with CBSV. By integrating transcriptomic data across tissues and time points, we demonstrate that rapid induction of immune genes in stem and root tissues is a hallmark of resistance. These findings lay the groundwork for functional characterization of key genes and the development of molecular markers for resistance breeding programs.

## 4. Materials and Methods

### 4.1 Cassava Genotypes, Virus Isolates, and Experimental Conditions

The cassava plants used in this study were obtained from the cassava collection of DSMZ Plant Virus Department at the Julius Kühn-Institute (JKI) in Braunschweig, comprising genotypes from the South American cassava germplasm accessions, provided by CIAT (Centro Internacional de Agricultura Tropical), Cali, Colombia and three cassava genotypes were selected: susceptible TMS 96/0304, root-restricted DSC 260, and resistant DSC 167, each of which exhibited a distinct response to CBSV infection (Sheat *et al*. 2019; Sheat *et al*. 2022; Sheat and Winter 2023). The virus isolate was obtained from infected TMS 96/0304 cassava plants with reference isolate (CBSV-Mo83) (DSMZ PV-0949, GenBank accession FN434436).

At JKI, the cassava plants of each genotype were cultivated and propagated in vitro and subsequently grown under identical conditions in a greenhouse. Plants were confirmed to be in a similar developmental and physiological state, propagated via tissue culture, and grown in a controlled greenhouse environment with temperatures between 26 °C and 30 °C, 12 hours light and 60 % humidity. The plants were potted in rectangular pots (12×12×20 cm) with Substrat GreenFibre from Klasmann-Deilmann GmbH. They were watered daily between 1 pm and 2 pm without any fertilizer. The workflow of the individual steps is shown in Figure 5.

**Figure 5:**
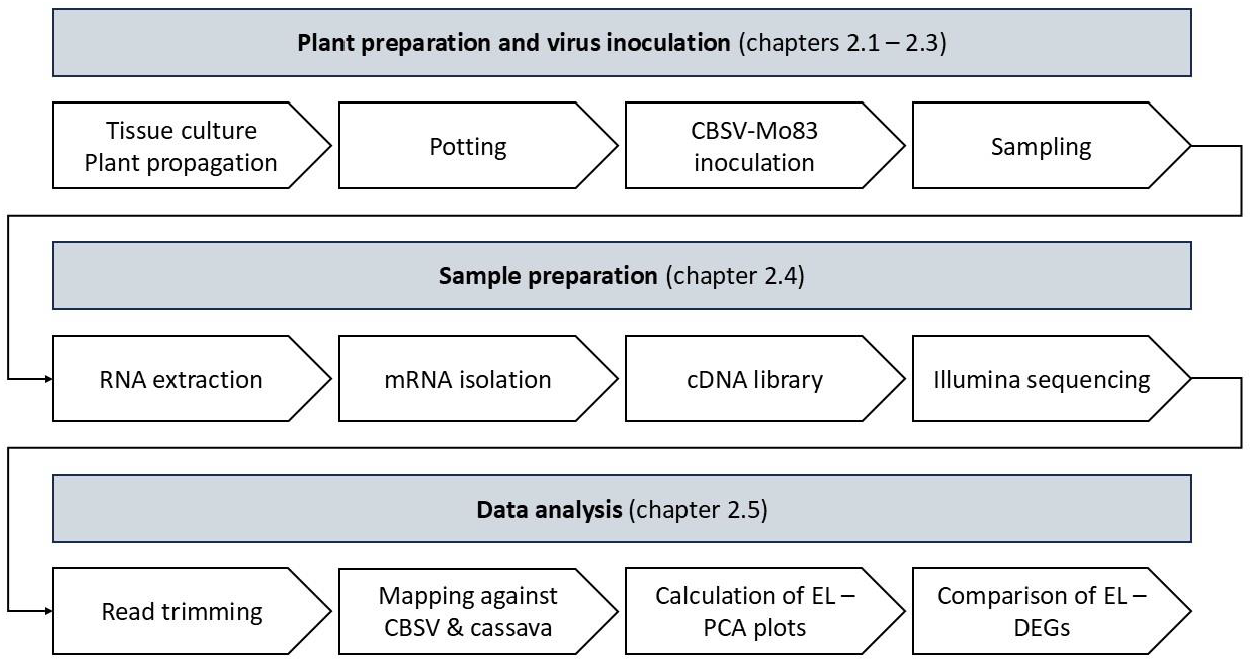
Sample preparation workflow.

### 4.2 Plant Infections

The cassava plants of TMS 96/0304 (susceptible), DSC 260 (root-restricted), and DSC 167 (resistant) were infected via bud grafting following the protocol outlined by Winter *et al*. 2010. Inoculation of the virus was conducted by grafting two axillary buds from the CBSV-infected TMS 96/0304 plant (Figure 5) onto the lower stem of each healthy plant; a total of 35 CBSV-infected TMS 96/0304 plants were used. A minimum of 15 healthy plants of TMS 96/0304, DSC 260, and DSC 167 were included for grafting the negative control groups, to account for transcriptional responses to inoculation wounding. The healthy plants of each genotype were grafted with buds from healthy plants of the same genotype.

In total, 54 plants per genotype were utilized for grafting and the bud grafting was performed between 8 am and 11 am in the middle of November. In addition to the 54 experimental plants, six backup plants were grafted for each genotype and condition (healthy or infected). The CBSV-infected backup plants of TMS 96/0304 were left in the greenhouse after the sampling for an extended period to observe whether symptoms would develop over time. This would serve to validate the efficiency of the virus inoculation via bud grafting.

### 4.3 Symptom Scoring and Sample Collection

Symptoms on leaves and stems were scored using a scoring system with S for virus symptoms and S0 for no symptoms, S+ for moderate symptoms, S++ for severe symptoms, S+++ for severe symptoms followed by plant death, and S? for inconspicuous symptoms (Sheat *et al*. 2019).

Samples of healthy and CBSV-infected plants were collected at three distinct time points: 1, 5, and 10 days after inoculation (DAI) (Figure 5). Samples were obtained from the stem below the bud grafting, the roots, the youngest fully expanded leaf (YL), and the first leaf from the bottom (OL). A total of nine plants were sampled per genotype, time point, plant part, and condition from infected and control plants. At each time point and for each condition, samples from three different plants were pooled together. A total of three replicates (A, B, and C) of each genotype were collected. The samples were frozen in liquid nitrogen and stored at -20 °C.

### 4.4 RNA Extraction

The flash frozen samples were ground using a tissue lyser (Qiagen TissueLyser LT, Germany) and RNA was extracted using the RNeasy Plant Mini Kit (Qiagen) (Figure 5), following the manufacturer’s protocol. Residual DNA was removed through a DNase I treatment performed according to the TAL protocol (Transcriptome Analysis Laboratory, Göttingen, Germany). The reaction mixture utilized for the DNase I treatment included 40 µl of extracted RNA, 20 µl of 10x incubation buffer, 4 µl of DNase I (10 U/µl), 2 µl of RNase OUT (40 U/µl), and 134 µl of RNase-free water. The mixture was incubated at 37 °C for 20 minutes. Following the incubation period, 100 µl of RNase-free water was added, followed by 300 µl of phenol/chloroform. The samples were vortexed and centrifuged at 15,304 x g for 2 minutes at room temperature.

The upper phase, which contained the RNA, was transferred to a new 1.5 ml tube. The precipitation of the RNA was initiated by the addition of 30 µl of 3 M sodium acetate (pH 4,8) and 300 µl of isopropyl alcohol. Following vortexing and a brief centrifugation, the tubes were stored at -20 °C for 30 minutes. Subsequently, the samples were subjected to centrifugation at 15,304 x g for 30 minutes at 4 °C. The supernatant was removed, and the RNA pellet was washed twice with 1 ml of 75 % ethanol, with centrifugation at 15,304 x g for 5 minutes at 4 °C each time. After the final wash, any residual ethanol was carefully removed, and the sediment was air-dried for approximately 10 minutes. The RNA pellet was resuspended in 30 µl of RNase-free water and mixed by pipetting. RNA quantities were verified using a Nanodrop™2000 (PEQLAB, Germany), with an additional quality check performed using an 1 % agarose gel. The RNA samples were stored at -80 °C until the library preparation for RNA-seq commenced.

### 4.5 cDNA Library and RNA-seq

The cDNA libraries were created using the Illumina Stranded mRNA Prep Kit, as supplied by Illumina (version June 2022). In a first step, the mRNA was purified using oligo(dT) magnetic beads which are provided in the Illumina kit. The enriched mRNA was then used to create the cDNA library for Illumina sequencing, in accordance with the manufacturer’s instructions in a cycler (Eppendorf SE, Germany). The final cDNA library samples were evaluated for integrity and quality using the Qubit DNA High Sensitivity assay (Invitrogen, Thermo Fisher Scientific) and the Agilent 2100 Bioanalyzer DNA 1000 High Sensitivity Kit (Agilent).

Later,the cDNA libraries were diluted to 2 nM to ensure uniform concentration for Illumina sequencing. Subsequently, 2 µl of each sample was transferred into a single 1.5 ml tube for sequencing. The Illumina NextSeq2000 with P3 and P2 FlowCells (paired-end 2×150nt with 200 cycles) was used for the sequencing of the samples.

### 4.7 Data Analysis

The initial analysis (Figure 5) of the data was conducted using the software Geneious Prime (version 2025.0.2). The “Trimming” function, utilising the R package BBDuk (https://sourceforge.net/projects/bbmap/ ; version 38.84), was implemented to remove the Illumina adapters and indexes from the raw reads. To confirm the detection of the virus, the reads were mapped to the CBSV-Mo83 genome sequence (FN434436.1, NCBI) using the Geneious RNA aligner from Geneious Prime (settings in Figure S4). Additionally, pairs and reads were aligned to the cassava reference genome sequence AM560-2, version 8 (GCF_001659605.2, NCBI), as well as to the reference subgenome sequences of cassava mitochondria (NC_045136.1, NCBI) and chloroplasts (NC_010433.1, NCBI). The paired-end reads and contigs were saved for subsequent analysis, with expression levels calculated in relation to the cassava reference genome sequence. A table, comprising transcripts per million (TPM) values for all 54 samples of one sampling area, was imported into R. PCA plots were created using the R tool ggplot2. Furthermore, to explain potential outliers in the dataset, rRNA checks were applied to check the remaining amount of rRNA in each sample. For the rRNA check the Python script rRNA_check.py (version v0.1) was used (https://github.com/bpucker/RNAseqQualCheck/blob/main/rRNA_check.py). DESeq2 (Love *et al*. 2014) was used to determine the expression levels between infected and healthy samples within the same genotype and time point, while the normalizing for sequencing depth and RNA composition was done using the median of ratios method. The resulting table, which shows the expression differences, was exported with columns for gene ID, protein ID, log2 ratio, and adjusted p-value. Genes with an adjusted p-value greater than 0.05 and log2 ratios between 1.0 and -1.0 were excluded in order to focus on significant differences. Bar charts were constructed to illustrate the total number of differentially expressed genes (DEGs) and their respective up- or down-regulation. The function of DEGs with log2 ratios greater than 1.0 or smaller than -1.0 was described using NCBI and compared in terms of their appearance and regulation. Pathway analyses were implemented by translating the protein IDs of the cassava samples into the corresponding gene IDs of *Arabidopsis thaliana* by using BLAST on NCBI against the *Arabidopsis thaliana* proteome. KEGG pathway analyses were applied by using The Arabidopsis Information Resource (TAIR) homologues to perform KEGG enrichment analyses in R.

## Author Contributions

For research articles with several authors, a short paragraph specifying their individual contributions must be provided. The following statements should be used “Conceptualization, X.X. and Y.Y.; methodology, X.X.; software, X.X.; validation, X.X., Y.Y. and Z.Z.; formal analysis, X.X.; investigation, X.X.; resources, X.X.; data curation, X.X.; writing—original draft preparation, X.X.; writing—review and editing, X.X.; visualization, X.X.; supervision, X.X.; project administration, X.X.; funding acquisition, Y.Y. All authors have read and agreed to the published version of the manuscript.” Please turn to the [CRediT taxonomy](https://img.mdpi.org/data/contributor-role-instruction.pdf) for the term explanation. Authorship must be limited to those who have contributed substantially to the work reported.

## Acknowledgments

We are grateful for the assistance during the experiments provided by the team of the Plant Virus Department of the DSMZ. This work was supported by the BMBF-funded de.NBI Cloud within the German Network for Bioinformatics Infrastructure (de.NBI) (031A532B, 031A533A, 031A533B, 031A534A, 031A535A, 031A537A, 031A537B, 031A537C, 031A537D, 031A538A). We also want to thank Google Colab for help to create the heatmap and create the bar chart of the rRNA check.

## References

Alicai, T.; Omongo, C. A.; Maruthi, M. N.; Hillocks, R. J.; Baguma, Y.; Kawuki, R. et al. (2007): Re-emergence of Cassava Brown Streak Disease in Uganda. In: Plant disease 91 (1), S. 24–29. DOI: 10.1094/PD-91-0024.

Alves, A. A. C. (2002): Cassava botany and physiology. In: R.J. Hillocks und J.M. Thresh (Hg.): Cassava: biology, production and utilization. UK: CABI Publishing, S. 67–89.

Amuge, T.; Berger, D. K.; Katari, M. S.; Myburg, A. A.; Goldman, S. L.; Ferguson, M. E. (2017): A time series transcriptome analysis of cassava (Manihot esculenta Crantz) varieties challenged with Ugandan cassava brown streak virus. In: Scientific reports 7 (1), S. 9747. DOI: 10.1038/s41598-017-09617-z.

Anjanappa, Ravi B.; Mehta, Devang; Maruthi, M. N.; Kanju, Edward; Gruissem, Wilhelm; Vanderschuren, Hervé (2016): Characterization of Brown Streak Virus-Resistant Cassava. In: Molecular plant-microbe interactions : MPMI 29 (7), S. 527–534. DOI: 10.1094/MPMI-01-16-0027-R.

Anjanappa, Ravi B.; Mehta, Devang; Okoniewski Michal J.; Szabelska-Berȩsewicz, Alicja; Gruissem, Wilhelm; Vanderschuren, Hervé (2018): Molecular insights into Cassava brown streak virus susceptibility and resistance by profiling of the early host response. In: Molecular plant pathology 19 (2), S. 476–489. DOI: 10.1111/mpp.12565.

Chen, Letian; Hamada, Satoshi; Fujiwara, Masayuki; Zhu, Tingheng; Thao, Nguyen Phuong; Wong, Hann Ling et al. (2010): The Hop/Sti1-Hsp90 chaperone complex facilitates the maturation and transport of a PAMP receptor in rice innate immunity. In: Cell host & microbe 7 (3), S. 185–196. DOI: 10.1016/j.chom.2010.02.008.

Dodds, Peter N.; Rathjen John P. (2010): Plant immunity: towards an integrated view of plant-pathogen interactions. In: Nature reviews. Genetics 11 (8), S. 539–548. DOI: 10.1038/nrg2812.

Hillocks, R. J.; Jennings, D. L. (2003): Cassava brown streak disease: A review of present knowledge and research needs. In: International Journal of Pest Management 49 (3), S. 225–234. DOI: 10.1080/0967087031000101061.

Hillocks, R. J.; Thresh, J. M.; Tomas, J.; Botao, M.; Macia, R.; Zavier, R. (2002): Cassava brown streak disease in northern Mozambique. In: International Journal of Pest Management 48 (3), S. 178–181. DOI: 10.1080/09670870110087376.

Howeler, R. H.; Lutaladio, NeBambi; Thomas, Graeme (2013): Save and grow. Cassava : a guide to sustainable production intensification. Rome: Food and Agriculture Organization of the United Nations.

Hu, Wei; Yan, Yan; Tie, Weiwei; Ding, Zehong; Wu, Chunlai; Ding, Xupo et al. (2018): Genome-Wide Analyses of Calcium Sensors Reveal Their Involvement in Drought Stress Response and Storage Roots Deterioration after Harvest in Cassava. In: Genes 9 (4). DOI: 10.3390/genes9040221.

Hubert, David A.; Tornero, Pablo; Belkhadir, Youssef; Krishna, Priti; Takahashi, Akira; Shirasu, Ken; Dangl Jeffery L. (2003): Cytosolic HSP90 associates with and modulates the Arabidopsis RPM1 disease resistance protein. In: The EMBO journal 22 (21), S. 5679–5689. DOI: 10.1093/emboj/cdg547.

Jones, Jonathan D. G.; Dangl Jeffery L. (2006): The plant immune system. In: Nature 444 (7117), S. 323–329. DOI: 10.1038/nature05286.

Kanzaki, H.; Saitoh, H.; Ito, A.; Fujisawa, S.; Kamoun, S.; Katou, S. et al. (2003): Cytosolic HSP90 and HSP70 are essential components of INF1-mediated hypersensitive response and non-host resistance to Pseudomonas cichorii in Nicotiana benthamiana. In: Molecular plant pathology 4 (5), S. 383–391. DOI: 10.1046/J.1364-3703.2003.00186.X.

Kaweesi, Tadeo; Kawuki, Robert; Kyaligonza, Vincent; Baguma, Yona; Tusiime, Geoffrey; Ferguson Morag E. (2014): Field evaluation of selected cassava genotypes for cassava brown streak disease based on symptom expression and virus load. In: Virology journal 11, S. 216. DOI: 10.1186/s12985-014-0216-x.

Love, Michael I.; Huber, Wolfgang; Anders, Simon (2014): Moderated estimation of fold change and dispersion for RNA-seq data with DESeq2. In: Genome biology 15 (12), S. 550. DOI: 10.1186/s13059-014-0550-8.

Maruthi, M. N.; Bouvaine, Sophie; Tufan Hale A.; Mohammed Ibrahim U.; Hillocks Rory J. (2014): Transcriptional response of virus-infected cassava and identification of putative sources of resistance for cassava brown streak disease. In: PloS one 9 (5), e96642. DOI: 10.1371/journal.pone.0096642.

Maruthi, M. N.; Hillocks, R. J.; Mtunda, K.; Raya, M. D.; Muhanna, M.; Kiozia, H. et al. (2005): Transmission of Cassava brown streak virus by Bemisia tabaci (Gennadius). In: Journal of Phytopathology 153 (5), S. 307–312. DOI: 10.1111/j.1439-0434.2005.00974.x.

Monger, W. A.; Seal, S.; Cotton, S.; Foster, G. D. (2001): Identification of different isolates of Cassava brown streak virus and development of a diagnostic test. In: Plant Pathology 50 (6), S. 768–775. DOI: 10.1046/j.1365-3059.2001.00647.x.

Munganyinka, Esperance; Margaria, Paolo; Sheat, Samar; Ateka Elijah M.; Tairo, Fred; Ndunguru, Joseph; Winter, Stephan (2018): Localization of cassava brown streak virus in Nicotiana rustica and cassava Manihot esculenta (Crantz) using RNAscope® in situ hybridization. In: Virology journal 15 (1), S. 128. DOI: 10.1186/s12985-018-1038-z.

Nichols, R. F. W. (1950): The Brown Streak Disease of Cassava. In: The East African Agricultural Journal 15 (3), S. 154– 160. DOI: 10.1080/03670074.1950.11664727.

Patil, Basavaprabhu L.; Legg James P.; Kanju, Edward; Fauquet Claude M. (2015): Cassava brown streak disease: a threat to food security in Africa. In: Journal of General Virology 96 (5), S. 956–968. DOI: 10.1099/jgv.0.000014.

Sheat, Samar (2020): Characterization of natural resistance in cassava against viruses causing the cassava brown streak disease. Dissertation. TU Braunschweig, Braunschweig, Germany. Plant Virus Department, DSMZ-Deutsche Sammlung von Mikroorganismen und Zelkulturen.

Sheat, Samar; Fuerholzner, Bettina; Stein, Beate; Winter, Stephan (2019): Resistance Against Cassava Brown Streak Viruses From Africa in Cassava Germplasm From South America. In: Frontiers in plant science 10, S. 567. DOI: 10.3389/fpls.2019.00567.

Sheat, Samar; Winter, Stephan (2023): Developing broad-spectrum resistance in cassava against viruses causing the cassava mosaic and the cassava brown streak diseases. In: Frontiers in plant science 14, S. 1042701. DOI: 10.3389/fpls.2023.1042701.

Sheat, Samar; Zhang, Xiaofei; Winter, Stephan (2022): High-Throughput Virus Screening in Crosses of South American and African Cassava Germplasm Reveals Broad-Spectrum Resistance against Viruses Causing Cassava Brown Streak Disease and Cassava Mosaic Virus Disease. In: Agronomy 12 (5), S. 1055. DOI: 10.3390/agronomy12051055.

Shirasu, Ken (2009): The HSP90-SGT1 chaperone complex for NLR immune sensors. In: Annual review of plant biology 60, S. 139–164. DOI: 10.1146/annurev.arplant.59.032607.092906.

Story, H. H. (1936): Virus Diseases of East African Plants: VI.-A Progress Report on Studies of the Disease of Cassava. In: The East African Agricultural Journal 2, S. 34–39.

Takabatake, Reona; Ando, Yuko; Seo, Shigemi; Katou, Shinpei; Tsuda, Shinya; Ohashi, Yuko; Mitsuhara, Ichiro (2007): MAP kinases function downstream of HSP90 and upstream of mitochondria in TMV resistance gene N-mediated hypersensitive cell death. In: Plant & cell physiology 48 (3), S. 498–510. DOI: 10.1093/pcp/pcm021.

Thresh, J. M. (2006): Control of tropical plant virus diseases. In: Advances in virus research 67, S. 245–295. DOI: 10.1016/S0065-3527(06)67007-3.

Tompa, Peter; Csermely, Peter (2004): The role of structural disorder in the function of RNA and protein chaperones. In: FASEB journal : official publication of the Federation of American Societies for Experimental Biology 18 (11), S. 1169–1175. DOI: 10.1096/fj.04-1584rev.

Winter, Stephan; Koerbler, Marianne; Stein, Beate; Pietruszka, Agnes; Paape, Martina; Butgereitt, Anja (2010): Analysis of cassava brown streak viruses reveals the presence of distinct virus species causing cassava brown streak disease in East Africa. In: Journal of General Virology 91 (Pt 5), S. 1365–1372. DOI: 10.1099/vir.0.014688-0.

Zhao, Yanxiao; He, Yong; Chen, Xinyue; Li, Ninghong; Yang, Tongqing; Hu, Tingting et al. (2024): Different viral effectors hijack TCP17, a key transcription factor for host Auxin synthesis, to promote viral infection. In: PLoS pathogens 20 (8), e1012510. DOI: 10.1371/journal.ppat.1012510.

